# Meditators probably show increased behaviour-monitoring related neural activity

**DOI:** 10.1101/2022.07.07.499152

**Authors:** Neil W Bailey, Harry Geddes, Isabella Zannettino, Gregory Humble, Jake Payne, Oliver Baell, Melanie Emonson, Sung Wook Chung, Aron T Hill, Nigel Rogasch, Jakob Hohwy, Paul B Fitzgerald

## Abstract

**Objectives:** Mindfulness meditation is associated with better attention function. Performance monitoring and error-processing are important aspects of attention. We investigated whether experienced meditators showed different neural activity related to performance monitoring and error-processing. Previous research has produced inconsistent results. This study used more rigorous analyses and a larger sample to resolve the inconsistencies.

**Methods:** We used electroencephalography (EEG) to measure the error-related negativity (ERN) and error positivity (Pe) following correct and incorrect responses to a Go/Nogo task from 27 experienced meditators and 27 non-meditators.

**Results:** No differences were found in the ERN (all *p* > 0.05). Meditators showed larger global field potentials (GFP) in the Pe after both correct responses and errors, indicating stronger neural responses (*p* = 0.0190, FDR-p = 0.152, np^2^ = 0.0951, BFincl = 2.691). This effect did not pass multiple comparison controls. However, single electrode analysis of the Pe did pass multiple comparison controls (*p* = 0.002, FDR-p = 0.016, np^2^ = 0.133, BFincl = 220.659). Meditators also showed a significantly larger Pe GFP for errors only, which would have passed multiple comparison controls, but was not a primary analysis (p = 0.0028, np^2^ = 0.1493, BF10 = 9.999).

**Conclusions:** Meditation may strengthen neural responses related to performance monitoring (measured by the Pe), but not specifically to error monitoring (although measurements of the Pe after errors may be more sensitive to group differences). However, only the single electrode analysis passed multiple comparison controls, while analysis including all electrodes did not, so this conclusion remains tentative.

## Introduction

Mindfulness meditation is associated with both cognitive and mental health benefits (Galante et al., 2021, Khoury et al., 2013, Gu et al., 2015, Gill et al., 2020, Im et al., 2021). Meta-analytic evidence suggests that mindfulness enhances attention, self-regulation, and executive function (Sumantry & Stewart, 2021; Jha et al., 2007; Tang et al., 2007; Xue et al., 2011, Im et al., 2021). These effects are perhaps unsurprising since the practice of mindfulness includes focusing attention on the present experience and redirecting attention back to the present experience when attention wanders - an activity that requires attention, self-regulation and executive function (Cardaciotto et al., 2008, Larson et al., 2013; Teper & Inzlicht, 2012). Functional brain imaging has shown increased activity in areas associated with these functions, such as the anterior cingulate cortex (ACC), dorsolateral prefrontal cortex (DLPFC), and insula (Allen et al., 2012; Tang et al., 2010; Tomasino & Fabbro, 2016; Xue et al., 2011, Falcone & Jerram, 2018). These increases in mindful attention have been found to underpin the mental health benefits of mindfulness practice (Gu et al., 2015).

Recent theoretical perspectives have attempted to provide a deeper explanation for the link between mindfulness, attention, and improved mental health by using a predictive coding theory of brain function (Manjalay et al., 2020, Verdonk & Trousselard, 2021, Lutz et al., 2019, Laukkonen & Slagter, 2021). The predictive coding theory suggests that because the brain must interact with the environment but only has access to the environment indirectly (through its sensory apparatus), the brain constructs a Bayesian model of its environment (Hohwy, 2012, Friston, 2010). This model is constituted of prior beliefs, predictions about the environment and an individual’s place within it, which are based on the individual’s actions (Hohwy, 2012, Friston, 2010). These priors are constantly updated by incoming sensory evidence (information from the body and environment), which is conceived of as prediction error weighted according to its expected precision, so that sensory evidence that is expected to be more precise attains more neural gain and has a stronger role in belief updating. The predictive coding theory suggests the main objective of brain function is to reduce prediction error – the mismatch between the prior beliefs and sensory evidence – over the long-term average. Prediction error minimization is achieved by both updating prior beliefs based on sensory evidence and making active inferences (altering the environment to match prior beliefs) (Hohwy, 2012, Friston, 2010).

Within the predictive coding theory, researchers have theorized that mindfulness practice might: 1) increase the precision of sensory evidence as a result of training in attending to sensations, which enhances synaptic gain of neurons processing prediction errors (Manjalay et al., 2020, Verdonk & Trousselard, 2021, Lutz et al., 2019, Laukkonen & Slagter, 2021); 2) increase the control over the selection of sensations for increased precision (Manjalay et al., 2020, Verdonk & Trousselard, 2021, Lutz et al., 2019, Laukkonen and Slagter, 2021); 3) loosen the precision with which prior beliefs are held, such that posterior evidence does not produce as much prediction error because posterior evidence is more commonly within the expected range of the priors (Manjalay et al., 2020, Laukkonen & Slagter, 2021); or alternatively, 4) increase the accuracy and precision of prior beliefs as a result of the long term increase in signal enhancement of the sensory evidence, such that priors are closer to posteriors and as such create less prediction error (Verdonk & Trousselard, 2021). Another four theories about how mindfulness affects the parameters of the predictive coding theory are described in the supplementary materials. This plethora of theories highlights one of the issues with explanations within the predictive coding theory: without experimental evidence, modulation in any of a wide array of predictive coding parameters could reflect a theoretical explanation for the effect of any given intervention. As such, empirical testing of potential differences in the parameters within the predictive coding theory is necessary to further our understanding of the mechanisms of mindfulness meditation.

One important concept for both attentional function and predictive coding accounts of the brain is performance monitoring. Cognition underlying error processing involves detecting errors in performance and adjusting cognitive resources to optimise performance. Error processing is integral to goal directed behaviour (Maurer et al., 2019), and behavioural errors are necessarily the result of predictive coding errors (an erroneous commission response on a response inhibition task reflects a mismatch between the prior expectation of perceiving a stimulus that is associated with a response, and subsequent evidence of a stimulus associated with withholding a response). As such, examining neural activity time locked to the commission of an error in mindfulness meditators is informative about the effects of mindfulness meditation on attention, as well as the effects of mindfulness meditation on the parameters of predictive coding models of the brain. Previous research has indicated that error processing in these types of tasks relies on the neural activity in the ACC (Larson et al., 2013). Predictive coding accounts have also suggested the ACC is a primary hub in projecting representations of bottom-up prediction errors to the DLPFC, which enables the modulation of predictions within the DLPFC to enable adaptive behaviour (Alexander & Brown, 2019). As such, measuring neural responses to errors may provide support for certain conjectures of how mindfulness affects the brain’s predictive coding function. In particular, an increase in the amplitude of neural responses to errors would support models suggesting mindfulness was associated with increased precision of sensory evidence or increased control over the selection of sensations for increased precision – models 1 and 2 from the list in the previous paragraph.

Given this background, it is no surprise that mindfulness mediation has been proposed to enhance error processing, and that studies have examined how mindfulness affects the neural correlates of error-processing using electroencephalography (EEG). Two event-related potentials (ERPs) related to error processing have been identified: the error related negativity (ERN) and error positivity (Pe) (Dehaene et al., 1994). The ERN is a negative potential produced by the ACC, which occurs within 100ms of error commission (Dehaene et al., 1994; Falkenstein et al., 2000; Larson et al., 2013). The Pe is a positive potential generated by the cingulate cortex and insula that occurs approximately 200 to 400ms after error commission (Herrmann et al., 2004, O’ Connell et al., 2007, Ullsperger et al., 2010). There is some ambiguity about the exact functional significance of the ERN and Pe. One prominent theory of ERN generation is the conflict monitoring theory. According to this theory, the ERN is generated by neural activity related to the processing of a conflict between the commission of an error response and the desired correct response (Larson et al., 2014). In tasks that rely on executive function, attention, and working memory, larger ERN amplitudes are found in contexts that generate more conflict but are also related to better performance (Larson & Clayson, 2011). The Pe is thought to reflect conscious recognition of errors and is modulated by level of attention, arousal, motivation, and affective response to an error (Falkenstein et al., 2000; Larson et al., 2013; Larson & Clayson, 2011). The Pe has also been implicated as a putative neural marker for the accumulated amount of evidence that participants have access to in order to decide whether they have committed an error (Steinhauser & Yeung, 2010). Larger Pe amplitudes are related to increased motivation and conscious processing of errors (Larson & Clayson, 2011).

To date, research investigating the impact of a mindfulness meditation intervention on the ERN and Pe has produced conflicting results, despite eleven published studies on the topic.

Three studies comparing a mindfulness practice condition to a control condition have shown increased ERN amplitudes (Fissler et al., 2017, Pozuelos et al., 2019, Saunders et al., 2016), but four have shown no differences (Eichel et al., 2020, Larson et al., 2013, Lin et al., 2019, Rodeback et al., 2020), and one further has shown a decreased ERN (Schoenberg et al., 2014). Similarly, two studies of the Pe have reported increased amplitudes in the mindfulness condition (Lin et al., 2019, Rodeback et al., 2020), two studies reported an increase in Pe amplitude in the mindfulness group, but also an increase in the waitlist or relaxation control groups (Schoenberg et al., 2014, Eichel & Stahl, 2020), two studies reported no difference (Bing-Canar et al., 2016, Smart & Segalowitz, 2017), and one study has reported a reduction in Pe amplitude in the mindfulness group (Larson et al., 2013).

The inconsistent and mixed pattern of results reported across the interventional studies conducted to date might be related to the small amount of mindfulness experience provided to the participants in these studies by the interventions. A recent meta-analysis has shown that neural changes from mindfulness practice relate to the amount of mindfulness experience (Falcone & Jerram, 2018), so studies of long-term meditators might be more likely to detect effects for error processing. However, experimental study designs investigating long-term meditation practice are prohibitively difficult to implement. Noting this as a limitation, three published studies have examined error processing in long-term meditators in cross-sectional study designs. However, the results of these studies are also inconsistent. In the first investigation of experienced meditators, Teper & Inzlicht (2012) observed larger ERN amplitudes in experienced meditators (with on average 3.19 years of meditation practice) compared to non-meditators, but no differences in Pe amplitudes. In a similar study, Andreu et al., (2017) observed a larger ERN, as well as a trend towards smaller Pe amplitudes (in both error and correct responses) in a sample of Vipassana meditators with an average of 5.1 years of experience compared to an athlete control group (with matched practice time). However, in a study that examined the ERN and Pe in experienced meditators, with the meditation group averaging 8.7 years of meditation experience (Bailey et al., 2019) a Bayesian statistical approach showed evidence against differences in both the ERN and Pe in these experienced meditators compared to controls.

These inconsistencies could also be related to methodological issues in analysing the ERN and Pe. Several EEG data processing steps can result in false positive or false negative results. These include: 1) the selection of electrodes for analysis - because ERPs are dipolar with negative and positive voltage peaks depending on the generating brain region, analyses focused on different electrodes has the potential to completely reverse an effect, 2) the selection of time windows for analysis - brain activity can be differentially modulated between groups only at specific timepoints, and arbitrary time window selection can increase false positive results (Kilner, 2013), and 3) the use of a subtraction baseline correction method. Baseline correction is intended to correct for slow drifts in EEG data, but the subtraction method has recently been shown to transpose a mirror image of the distribution of activity during the baseline period onto the active period, while ironically also decreasing the signal to noise ratio (Alday, 2019). A more detailed explanation of these potential confounds can be found in the supplementary materials.

Our previous study was the first to examine error processing in meditators using methods that eliminate the possibility of bias from experimenter selection of electrodes or time windows from influencing results, addressing issues 1 and 2 listed above [BLINDED FOR REVIEW]. Because our null results conflicted with the previous literature, we here sought to replicate our previous study with a larger sample size and with the application of a more robust regression baseline correction method (Alday, 2019). This regression baseline correction has been demonstrated to adequately correct for slow drifts in the EEG data without transposing the mirror image of the baseline activity into the active period, removing this potential confound, and without decreasing the signal to noise ratio (Alday, 2019), addressing issue 3 listed above. Additionally, while most previous research has focused specifically on error processing, it is possible that meditation produces a general difference in performance monitoring, rather than a specific error-processing-related difference (as suggested by Andreu’s (2017) results). To explore this, we examined neural activity across both error and correct trial types. Based on initial findings from Teper and Inzlicht (2012) and Andreu et al., (2017), our alternative hypothesis was that meditators would have increased ERN amplitudes to both correct and error responses compared to controls. We also nominated an alternative hypothesis that meditators would show larger Pe amplitudes to both correct and error responses than non-meditators, in line with a slight majority of interventional studies.

However, it should be noted that given our previous null result study and the inconsistency in the literature, our primary hypothesis was that we would replicate the null result. As such, we planned to explore potential positive results fully, to provide a fair evaluation of our research question, and so that our conclusions would be protected against potential biases.

## Method

### Participants

Thirty-nine meditators and 37 healthy non-meditators were recruited via community advertising at universities, meditation organisations, and social media. Of these participants, 57 participants (30 meditators and 27 controls) provided enough artifact free error response EEG epochs for analysis (details below). Data were collected by students trained in EEG data recording and supervised by experienced researchers. The study had no direct funding for personnel, so no a priori power analysis was performed. As such, the intended sample size was simply “larger than our previous study, and as many as possible” within the resource and time restrictions. The final sample for the two groups significantly differed in age, so the three oldest meditators were excluded prior to analysis, so that the groups did not significantly differ in any demographic variables (all *p* > 0.10, note that in order to further reduce the influence of the potential confound, analyses were performed controlling for age and years of education as covariates). The final data set comprised a total of 54 participants (27 meditators and 27 controls). To be considered an experienced mediator, participants were required to have practiced meditation for a minimum of two years. They also had to practice meditation for a minimum of two hours per week over the last three months. The meditation sample had an average of 7.57 years’ meditation experience (SD = 7.04), and an average of 7.98 hours of practice per week (SD = 5.82). To be included in the study, we required that meditators reported practicing in a way that is consistent with the following definition of mindfulness: “paying attention in a particular way: on purpose, in the present moment, and nonjudgmentally” (Kabat-Zinn, 1994), and that the meditator’s practice was focused on sensations of the breath or body. Trained meditation researchers screened and interviewed participants to ensure that their meditation practices were consistent with the criteria. Non-meditating control participants were eligible to participate if they had less than two hours meditation experience across their lifetime. Any uncertainties in screening were resolved by two researchers, including the principal researcher (BLINDED FOR REVIEW).

Participants were ineligible to participate if they had previous or current neurological or mental illness. Participants were also ineligible if they were currently taking psychoactive medication. All participants were interviewed with the MINI International Neuropsychiatric Interview (MINI) DSM-IV (Hergueta et al., 1998). Potential participants who met the criteria for psychiatric illness were excluded. The Beck Anxiety Inventory (BAI) and Beck Depression Inventory II (BDI-II) (Beck et al., 1996) were also administered. Any participants who scored above the moderate range in the BAI (>25) or greater than the mild range in the BDI-II (>19) were excluded. All included participants were aged between 19 and 61 years.

Prior to EEG recording, participants provided their age, gender, years of education, and mediation experience (total years of practice, frequency of practice, and amount of time spent practicing). In addition to the BAI and BDI-II, participants also completed the Five Facets of Mindfulness Questionnaire (FFMQ). These measures are summarised in Table 1. Ethics approval was provided by the Ethics Committees of Monash University and Alfred Hospital. All participants provided written informed consent prior to participation in the study.

**Table 1.**
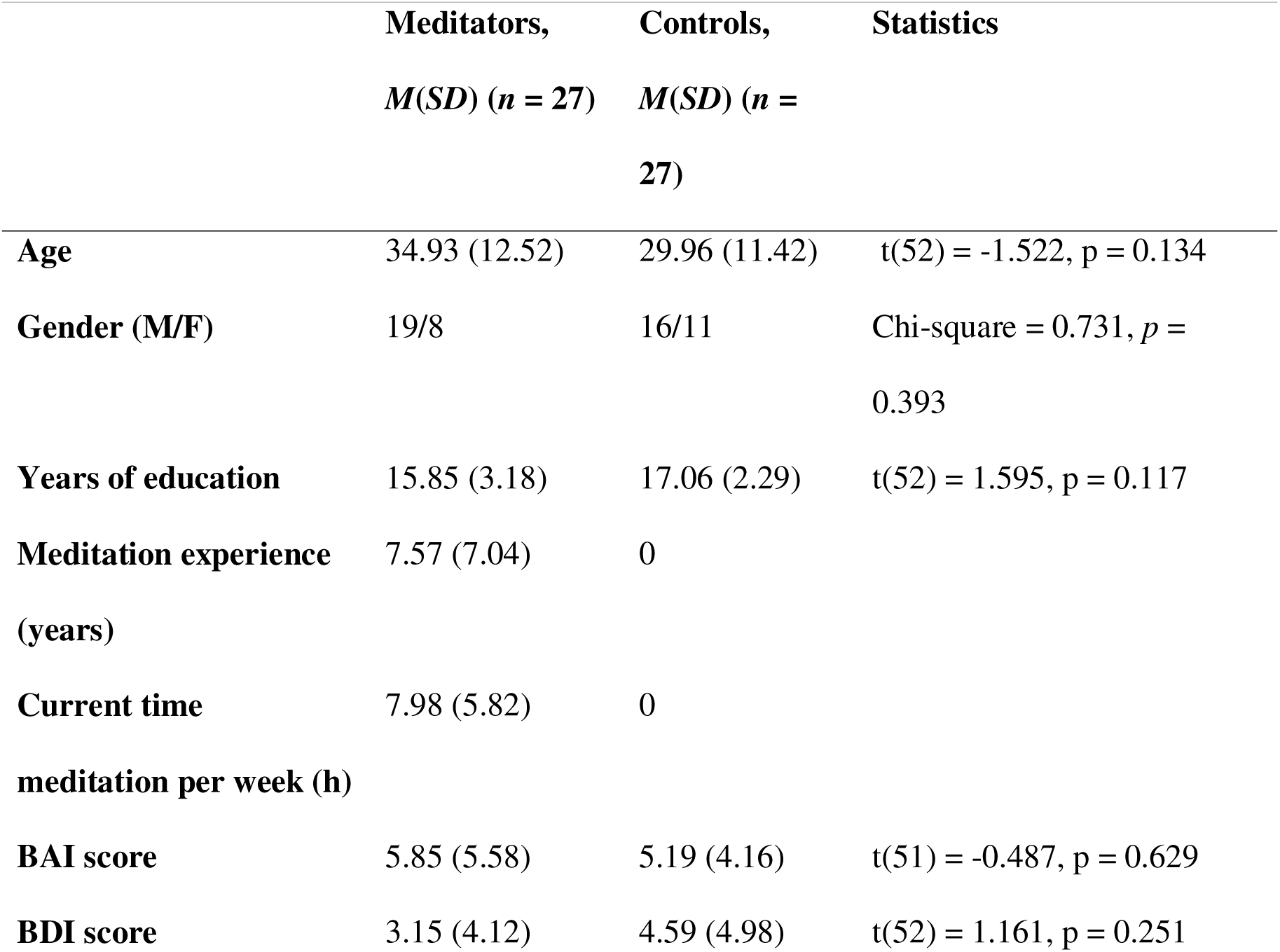
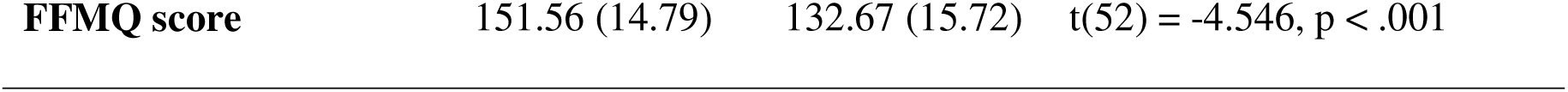
Participant demographic data. Note that BAI data was missing for one control participant.

### Procedure

Sixty-four channel EEG was recorded while participants performed a Go/Nogo task (Neuroscan, Ag/AgCl Quick-Cap through a SynAmps 2 amplifier [Compumedics, Melbourne, Australia]). Electrodes were referenced to an electrode between Cz and CPz, impedances were kept below 5kΩ, and EEG was sampled at 1000Hz with online bandpass filters from 0.1 to 100Hz. The EEG was recorded while participants performed a Go/Nogo task with simplified emotional faces (stimuli were identical to BLINDED FOR REVIEW). The Go/Nogo task is commonly used to elicit ERN and Pe components and has been shown to have high reliability (alpha = 0.75, the highest out of the Go/Nogo, Flanker and Stroop tasks tested) (Clayson, 2019). The task included four separate blocks. The first two blocks were an easy version of the task, each with 50 happy faces and 50 sad faces. One block required participants to respond (Go) to happy faces and withhold response (Nogo) to sad faces. Participants who responded to happy faces in the first block responded to sad faces in the second block, and vice versa. The stimulus-response pairing was counter-balanced across participants, so half of the participants in each group responded to happy faces in the first block and sad faces in the second block, and the other half responded to sad faces in the first block then happy faces in the second block. Following this, two harder blocks were presented, each with 50 Nogo trials and 150 Go trials (again with the stimulus-response pairing counterbalanced). Each stimulus was presented for 250ms with an intertrial interval of 900ms (with a 50ms jitter). EEG data were pre-processed using the RELAX pipeline, which has demonstrated optimal cleaning of artifacts and preservation of ERP signals compared to other cleaning approaches (Bailey et al., 2022a, 2022b). This cleaning pipeline filtered the data from 0.25 to 80Hz with a notch filter from 47 to 53Hz, applied both Multiple Wiener Filters (MWF) and wavelet enhanced independent component analysis (wICA) to identify and remove muscle, eye movement and blink and drift artifacts from the data, and additionally the wICA reduced line noise, heartbeat and other artifacts. Data were re-referenced to the average of all electrodes. Full details of the cleaning method are reported in the supplementary materials.

Following the cleaning of continuous files, data were epoched around correct and error responses from -400 to 800ms. Each error response (following a Nogo trial) was matched in reaction time to a correct response with the closest available reaction time (from a Go trial that presented a face of the same emotion) so that an equal number of error and correct responses were epoched for analysis, and these responses were matched for both condition and reaction time. Epochs with voltages exceeding +/-60 μV at any electrode were rejected, as were epochs containing improbable voltage distributions or kurtosis values >5SD from the mean in any single electrode or more than 3SD from the mean over all electrodes. Data were then baseline corrected by regressing out the average of the -400 to -100ms period from each epoch (timelocked to the response) using the fieldtrip function ‘ft_regressconfounds’ for each electrode and each participant separately, with the condition of each epoch (correct or error response) included in the regression model (but not rejected) to correct for potential voltage drift but still preserve any experimental effects (Bailey et al., 2022a, 2022b).

Participants who had less than six error related epochs remaining for analysis were excluded at this stage. We ensured our data provided reliable analysis by using the ERA Reliability Analysis Toolbox v0.5.3 (Clayson & Miller, 2017a, 2017b). The ERA Toolbox showed our dataset provided >0.90 dependability for both correct and error trials within both the averaged ERN and Pe windows. Further dependability details can be found in the supplementary materials.

### Data Analysis

#### Primary comparisons

Primary EEG data statistical comparisons were conducted using the Randomisation Graphical User Interface (RAGU) toolbox (Koenig et al., 2011). Unlike approaches that focus on specific electrodes and time windows for analyses, RAGU is a reference free approach that compares scalp field differences across all electrodes and time points with permutation statistics that are robust to the assumptions of traditional parametric statistics. As such, RAGU minimises need for a priori choices of time windows or electrodes, which can bias analyses. RAGU is also robust against the violation of the assumptions of traditional parametric statistics (Koenig et al., 2011).

RAGU also allows for separate comparisons of the overall neural response strength (using the global field power [GFP] test) and the distribution of neural activity across electrodes (with the Topographic Analysis of Variance [TANOVA]). We used the TANOVA to assess whether the distribution of neural activity differed between groups or conditions without the influence of overall neural response strength by normalising the overall amplitude of the neural response, so that all participants and conditions have a GFP = 1 prior to the TANOVA (using the recommended L2 normalisation). Additionally, prior to the TANOVA, a topographical consistency test (TCT) was conducted to ensure consistent distribution of scalp activity within each group/condition. The TCT is analogous to a single sample t-test, and assessed whether the signal within a condition or group significantly differed from 0 (in which case the group/condition demonstrated a consistent distribution of neural activity following a response).

The GFP and TANOVA tests were applied in a repeated measures ANOVA design, comparing 2 group (meditators vs controls) × 2 condition (corrects vs errors) in data from - 400 to 700ms around correct/error responses. Tests were conducted with 5000 permutations and an alpha of *p* = 0.05. To control for multiple comparisons across time, the global duration control was used, which ensures any significant effects last longer than 95% of the ‘significant effects’ within the randomly shuffled data. When significant effects were detected, we averaged data within the significant period, and report p-values, effect sizes, and Bayesian statistical evidence for the alternative hypothesis for these comparisons (this approach maximises effect size estimation by focusing only on the significant period). We also analysed data averaged across windows from typical ERN and Pe windows (50 to 150ms following the response and 200 to 400ms following the response respectively, this approach reduces effect size estimation by including potentially non-significant time periods). We extracted the average GFP for significant periods and windows of interest for Bayesian statistical analysis using JASP (Love et al., 2019) (however, note that Bayesian approaches are currently not available to replicate the TANOVA test of differences in the distribution of activity). Lastly, we performed experiment-wise multiple comparison controls using the Benjamini-Hochberg (1995) false discovery rate (FDR-p) for the traditional ERN and Pe time window of interest analyses, including all group main effects and interactions involving group in the multiple comparison controls for both the ERN and Pe across both the GFP and TANOVA tests (8 tests).

#### Replication Comparisons – Single Electrode Analysis

In addition to the whole scalp analyses conducted to test our primary hypotheses, we performed a traditional electrodes-of-interest analysis of the average ERN and Pe periods using five midline electrodes. We implemented a repeated measures ANOVA with the following design: group (meditator/control) x condition (correct/error) x electrode (Fz, FCz, Cz, CPz and Pz). Full details of this analysis are reported in the supplementary materials.

Because these analyses overlap with the primary analyses performed in RAGU, we performed experiment-wise multiple comparison controls using FDR-p separately for these analyses, including all group main effects and interactions involving group in the controls for both the ERN and Pe (8 tests).

Lastly, because our results differed from our previous study, we performed a re-analysis of the data from that study focused on error trials (which provide the largest signal), using the same methods as in the current study. We report this analysis, and an analysis of the combined data from the current study and our previous study in our supplementary materials. No analyses were performed on behavioural data, as the behavioural comparisons from the full dataset are planned for a study in preparation which will examine stimulus locked EEG activity from the Go Nogo task.

## Results

### Demographic and Behavioural Data

Meditators and controls did not significantly differ in age, BDI, or BAI (all *p* > 0.10). However, as expected, meditators reported higher FFMQ scores than controls (meditators = 151.56 (14.79), controls = 132.67 (15.72)), *t*(52) = -4.546, *p* < .001. Means suggested the behavioural performance was similar across groups (Table 2).

**Table 2.** Participant Behavioural Data and number of accepted epochs.

|  | **Meditators** | **Controls** |
| --- | --- | --- |
|  | **M (SD)** | **M (SD)** |
| **Number of accepted correct epochs** | 28.74 (16.05) | 33.86 (19.31) |
| **Number of accepted error epochs** | 27.59 (16.02) | 33.44 (20.33) |
| **Percent correct equal Go trials** | 95.23 (12.52) | 96.00 (9.28) |
| **Percent correct equal Nogo trials** | 92.08 (12.90) | 91.11 (12.00) |
| **Percent correct frequent Go trials** | 95.85 (9.60) | 97.47 (2.74) |
| **Percent correct infrequent Nogo trials** | 70.43 (16.92) | 67.74 (17.71) |
| **Overall Correct Go RT** | 374.01 (42.72) | 366.45 (39.26) |
| **Overall Correct Hard Go RT** | 336.97 (52.83) | 320.85 (39.39) |
| **Error Nogo RT** | 299.22 (65.36) | 282.18 (59.29) |
| **Hard Error Nogo RT** | 259.13 (30.41) | 255.59 (36.92) |
| **Matched correct equal Go RT** | 314.38 (71.44) | 286.85 (53.85) |
| **Matched correct frequent Go RT** | 262.63 (33.68) | 257.71 (37.58) |

### Primary Comparisons

#### ERN and Pe Global Field Potential Tests

The GFP test showed no significant main effect of group or interaction between group and response that lasted longer than global duration controls (67ms for the main effect of group, 39ms for the interaction between group and response). However, the meditator group showed intermittent periods with larger Pe amplitudes across multiple time periods spanning 235 to 373ms, none of which individually passed the global duration statistic, but the entire period would have passed global duration controls if it were not intermittent. Additionally, this period was in the Pe window of interest (Figure 1A). To protect against the possibility of accepting a null result (in line with the null results of our previous study, BLINDED FOR REVIEW) where there was suggestive evidence of a positive result, we explored this effect further. When averaging across this 235 to 373ms period, the main effect of group was significant, with the meditator group showing a higher amplitude GFP than controls across both conditions (*p* = 0.0186, np^2^ = 0.1066, BFincl = 2.697, see Figure 1D). Additionally, when averaged across the typically analysed Pe period (200 to 400ms), the main effect of group was significant with the meditator group showing a significantly higher amplitude GFP than controls across both conditions (*p* = 0.019, FDR-p = 0.1520, np^2^ = 0.0951, BFincl = 2.691) (however note that the difference did not pass our experiment-wise multiple comparison control). This may indicate that meditators have larger neural response amplitude (independent of the distribution of activity) than controls to both error and correct response trials during the Pe window. There was a trend towards an interaction between group and response when data was averaged within the 235 to 373ms period (p = 0.0738, np^2^ = 0.0622, BFincl = 1.257), however, this effect was not present when averaging across the typical Pe window (200 to 400ms, p = 0.12, FDR-p = 0.2496, np^2^ = 0.0535, BFincl = 1.300), and no interaction between group and response was present that passed duration controls. This suggests the potential difference in Pe amplitude was not specific to error responses (that the main effect of group was larger than the effect for a specific condition). In contrast, when data was averaged within the ERN window of interest (50 to 150ms), there was no significant difference between groups (p = 0.1248, FDR-p = 0.2496, np^2^ = 0.0453, BFexcl = 0.962) nor interaction between group and condition (p = 0.7028, FDR-p = 0.9371, np^2^ = 0.0030, BFexcl = 3.736).

**Figure 1.**
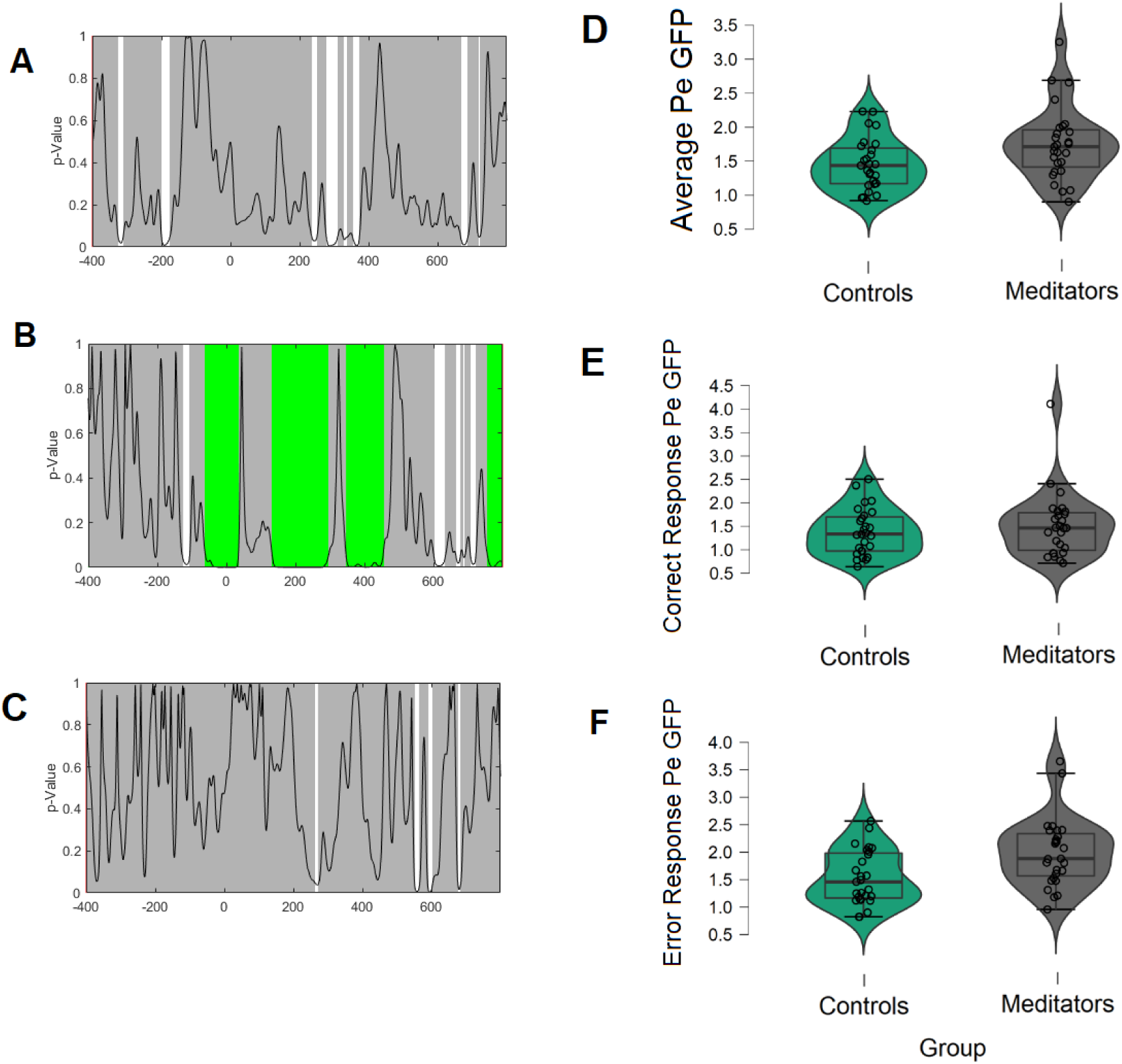
Significance p-graphs and violin plots for the GFP comparisons in the novel dataset. A: the p-map for the main effect of group. B: the p-map for the main effect of response condition. C: The p-map for the interaction between group and condition. For A-C, the black line reflects the *p* value, grey periods reflect no significant differences between groups, white periods reflect significant differences that did not survive duration multiple comparison controls. D: Averaged GFP values during the typical Pe window of interest 200 to 400ms for both responses averaged together. E: Averaged GFP values during the typical Pe window of interest 200 to 400ms for correct responses. F: Averaged GFP values during the typical Pe window of interest 200 to 400ms for error responses. Note that the significant main effect of group indicated that the groups differed in GFP when averaged across the two response conditions.

**Figure 2.**
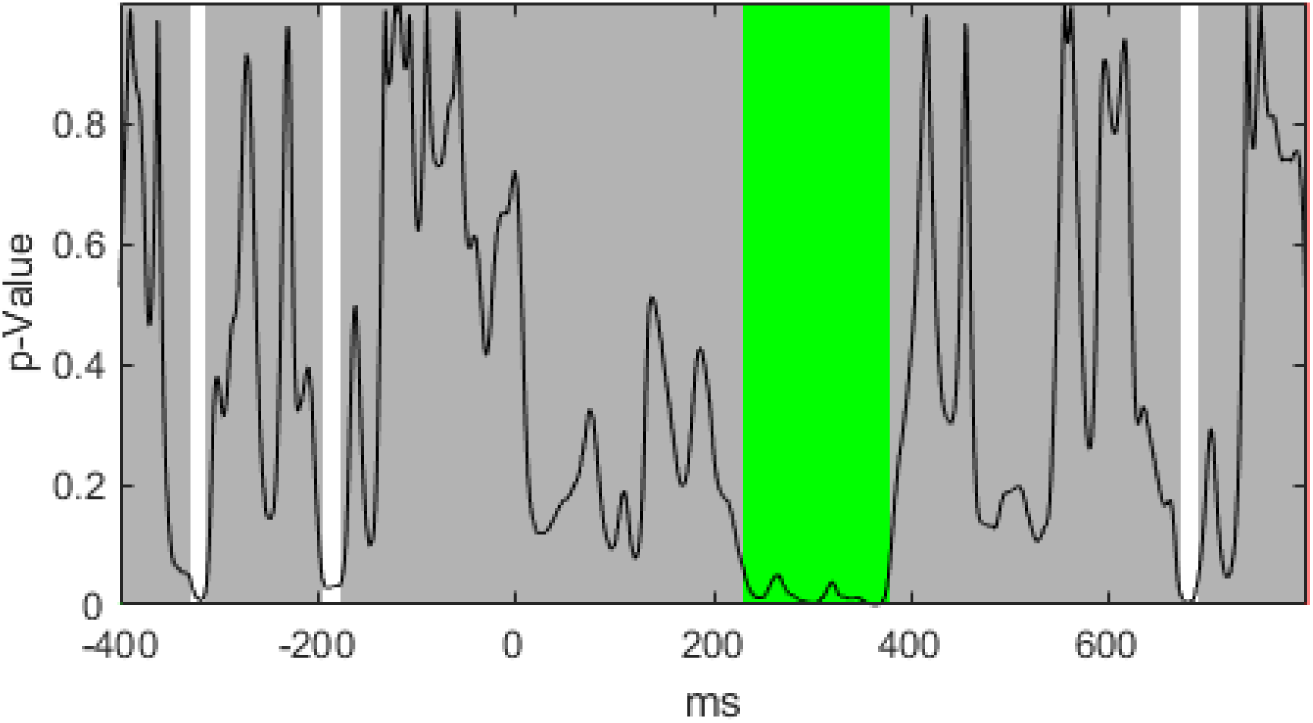
Significance p-graph for the GFP comparisons between groups when including error responses only.

Next, while the groups did not significantly differ in age or years of education, they were not directly matched. To ensure these potential confounds were not influencing our results, we performed the comparison in RAGU averaged across the Pe window of interest, with age and years of education regressed out of the analysis. Again, this comparison showed a significant main effect of group with meditators showing larger amplitudes (p = 0.034, np^2^ = 0.0840). We also performed a repeated measures ANCOVA comparison of the averaged Pe window using JASP, covarying for age and years of education. These results showed the main effect of group was reduced to a trend F(1,50) = 3.072, p = 0.086, np^2^ = 0.058, BFincl = 1.125, but that the interaction between group and response (correct/error) was significant F(1,50) = 7.849, p = 0.007, np^2^ = 0.136, BFincl = 3.914. Separate ANCOVA analyses for the error and correct trials indicated that this was due to a significant effect of group in the error response condition F(1,50) = 10.352, p = 0.002, np^2^ = 0.172, BFincl = 15.757, but not the correct response condition F(1,50) = 0.026, p = 0.873, np^2^ = 5.151e-4, BFexcl = 3.205.

It is worth noting that age was non-significantly negatively correlated with the error Pe GFP (r = -0.099, p = 0.478), so if age had affected our results, the difference between meditators and controls would have been reduced, making a null result more likely (as older participants showed smaller Pe GFP and the meditation group was older on average). Years of education was also not significantly correlated with Pe GFP (r = -0.178, p = 0.198). Additionally, age and years of education were not significant predictors in the ANCOVA, Age: F(1,50) = 0.243, p = 0.624, BFexcl = 3.183, Years of education: F(1,50) = 0.789, p = 0.379, BFexcl = 2.127.

Although the interaction between group and response was not significant in our primary analysis, it was significant when covarying for age. Additionally, error processing studies often focus only on the error related responses, and error ERPs typically generate more signal than correct ERPs. Given this information, and the significant interaction when covarying for age, as well as our aim to protect against potential biases towards a null result, we performed an exploratory analysis focused only on the error trials. This analysis showed a group main effect from 230 to 376ms, which lasted longer than the global duration control (57ms).

Averaged across this interval, the effect was significant with a large effect size (p = 0.0022, np^2^ = 0.1553, BF10 = 11.813). Averaged across the typical Pe window of interest (200 to 400ms), the effect remained significant (p = 0.0028, np^2^ = 0.1493, BF10 = 9.999). We also note here that if we had planned our analysis to focus on error trials only and included this result in our multiple comparison controls, the difference in error related Pe GFP would have passed our multiple comparison controls (FDR-p = 0.0176). It is also worth noting that while this effect was not significant in the data from our previous study (after re-processing using the current study’s methods), the pattern was in the same direction (p = 0.176, np^2^ = 0.0576, BF10 = 0.658, Figure S5). Additionally, the combined analysis across both the current and our previous dataset was significant (p = 0.019, np^2^ = 0.0576, BF10 = 2.607, or BF10 = 5.147 for a one-sided test, given the sample in our current study suggested this would be the pattern, Figure S7). However, it is also worth noting that the group by condition interaction in our primary analysis was not significant, so traditionally we would not have explored the error-related Pe in isolation.

#### ERN and Pe TANOVA

The TCT indicated mostly consistent distributions of neural activity within each condition for each group, including during the ERN and Pe windows of interest, suggesting it was valid to perform comparisons using the TANOVA (with the exception of a brief period around 120ms in the meditation error condition, and around 200ms for the correct condition for both groups, Figure S2). The TANOVA showed a significant main effect of condition across the entire epoch starting in the baseline period (-86ms), suggesting error and correct responses generated different distributions of neural activity across the entire period. Both the ERN and Pe showed the typical scalp distribution pattern (Figure S3). However, no significant main effect of group lasting longer than the global duration controls (51ms) was found in the TANOVA (very brief effects were present at 288 to 304ms and 449 to 461ms, see Figure S4). There was also no significant interaction between group and response (correct/error) that lasted longer than the global duration controls (39ms, all *p* > 0.05, global count p = 0.710). This result suggests that the two groups did not differ in the distribution of brain activity time locked to responses, nor did the distribution of brain activity interact between the groups and type of correct or error response. Averaged within the ERN window of interest, there was no significant main effect of group (p = 0.9508, FDR-p = 0.9646, np^2^ = 0.0052), nor an interaction between group and response (correct/error) (p = 0.6554, FDR-p = 0.9371, np^2^ = 0.0174). Averaged within the Pe window of interest, there was no significant main effect of group (p = 0.0826, FDR-p = 0.2496, np^2^ = 0.0367), nor an interaction between group and response (correct/error) (p = 0.9646, FDR-p = 0.9646, np^2^ = 0.0060).

### Replication Comparisons – Single Electrode Analysis

There was no significant main effect for the ERN group comparison, nor interactions between group and response (error/correct) or interactions between electrode, group and response (error/correct) (all *p* > 0.45, BFexcl for the main effect of group = 5.378, BFexcl for response * group = 7.465, and for the interaction between group * electrode and group * electrode * response type, BFexcl > 18). Averaged activity across the ERN windows at each electrode can be viewed in Figure S1 (and full statistics can be viewed in Table 3).

**Table 3.** Means, standard deviations, and full statistics for the Pe GFP and the electrode of interest analysis.

| Comparison | Meditators<br>M (SD) | Controls<br>M (SD) | Statistics |
| --- | --- | --- | --- |
| **GFP Error 235 to 373ms** | 1.968 (0.709) | 1.446 (0.525) | Group main effect: $p = 0.0186$, $np^2 = 0.1066$, $BFincl = 2.697$ |
| **GFP Correct 235 to 373ms** | 1.531 (0.695) | 1.401 (0.510) | Response x Group: $p = 0.0738$, $np^2 = 0.0622$, $BFincl = 1.257$ |
| **GFP Error 200 to 300ms** | 1.993 (0.622) | 1.533 (0.491) | Group main effect: $p = 0.0190$, $FDR-p = 0.152$, $np^2 = 0.0951$, $BFincl = 2.691$ |
| **GFP Correct 200 to 300ms** | 1.513 (0.688) | 1.382 (0.499) | Response x Group: $p = 0.12$, $FDR-p = 0.2496$, $np^2 = 0.0535$, $BFincl = 1.300$ |
| **ERN Error Fz** | -0.80 (2.39) | -0.90 (1.48) | Group main effect: |
| **FCz** | -1.56 (2.56) | -1.26 (1.94) | $F(1,52) = 0.095$, $p = 0.760$, $FDR-p = 0.869$, $np^2 = 3.513e-4$, $BFexcl = 5.378$ |
| **Cz** | -1.33 (1.98) | -0.90 (1.96) | Response x Group: |
| **CPz** | -0.49 (1.33) | -0.45 (1.51) | $F(1,52) = 0.025$, $p = 0.875$, $FDR-p = 0.875$, $np^2 = 4.808e-4$, $BFexcl = 7.465$ |
| **Pz** | 0.21 (1.54) | -0.21 (1.09) | Group x Electrode: |
| **ERN Correct Fz** | -0.59 (2.03) | 0.20 (1.43) | $F(1,52) = 0.702$, $p = 0.458$, $FDR-p = 0.734$, $np^2 = 0.013$, $BFexcl = 18.189$ |
| **FCz** | 0.45 (1.81) | 0.80 (1.50) | Response x Electrode x Group: |
| **Cz** | 1.22 (1.11) | 1.39 (1.23) | $F(1,52) = 0.752$, $p = 0.459$, $FDR-p = 0.734$, $np^2 = 0.014$, $BFexcl = 27.562$ |
| **CPz** | 1.27 (1.01) | 1.19 (0.89) |  |
| **Pz** | 0.90 (1.28) | 0.52 (0.93) |  |
| **Pe Error Fz** | 2.19 (1.94) | 1.29 (1.29) | Group main effect: |
| **FCz** | 2.67 (1.86) | 1.75 (1.22) | $F(1,52) = 2.848, p = 0.097, \text{FDR-}p = 0.388, \text{np}^2 = 0.052, \text{BF}_{\text{excl}} = 2.250$ |
| **Cz** | 1.95 (1.32) | 1.56 (1.02) | Response x Group: |
| **CPz** | 0.95 (0.97) | 1.04 (0.84) | $F(1,52) = 0.886, p = 0.351, \text{FDR-}p = 0.734, \text{np}^2 = 0.017, \text{BF}_{\text{excl}} = 2.354$ |
| **Pz** | 0.08 (1.00) | 0.42 (0.83) | Group x Electrode: |
| **Pe Correct Fz** | 1.27 (0.91) | 0.81 (1.02) | $F(1,52) = 7.969, p = 0.002, \text{FDR-}p = 0.016, \text{np}^2 = 0.133, \text{BF}_{\text{incl}} = 220.659^*$ |
| **FCz** | 1.00 (1.05) | 0.58 (1.06) | Response x Electrode x Group: |
| **Cz** | 0.11 (0.81) | 0.02 (0.89) | $F(1,52) = 0.344, p = 0.619, \text{FDR-}p = 0.825, \text{np}^2 = 0.007, \text{BF}_{\text{excl}} = 24.373$ |
| **CPz** | -0.74 (0.67) | -0.56 (0.71) |  |
| **Pz** | -1.22 (0.76) | -0.84 (0.72) |  |

For the Pe component, a significant interaction between group and electrode was present, with clear evidence for the alternative hypothesis F(1,52) = 7.969, *p* = 0.002, np^2^ = 0.133, BFincl = 220.659. This aligned with the GFP results, which indicated the meditation group showed larger overall neural response strength, so post-hoc exploration of the cause of the effect was undertaken. The post-hoc exploration indicated that the two groups differed at Fz (p-Holm = 0.006) and FCz (p-Holm = 0.012) but not at other electrodes (all p-Holm > 0.50). Additionally, within the control group, Fz, FCz and Cz showed more positive voltages than CPz and Pz (all p-Holm < 0.011) while showing no other differences (all p-Holm > 0.08). In contrast, the meditation group showed a larger differentiation of voltage for each electrode than the control group, with all electrodes differing from all other electrodes, except Fz and FCz (all p-Holm > 0.002 except for Fz compared to FCz, for which p-Holm = 1.0). The results for the single trial analysis of the Pe can be visualised in Figure 3. While our re- analysis of our previous data did not show this significant interaction, the pattern was in the same direction (F(1,40) = 0.951, p = 0.378, np^2^ = 0.023, Figure S6). Additionally, our analysis of the combined dataset still showed strong support for the interaction between group and electrode (F(1,94) = 4.291, p = 0.025, np^2^ = 0.044, BFincl = 11.115, Figure S8).

**Figure 3.**
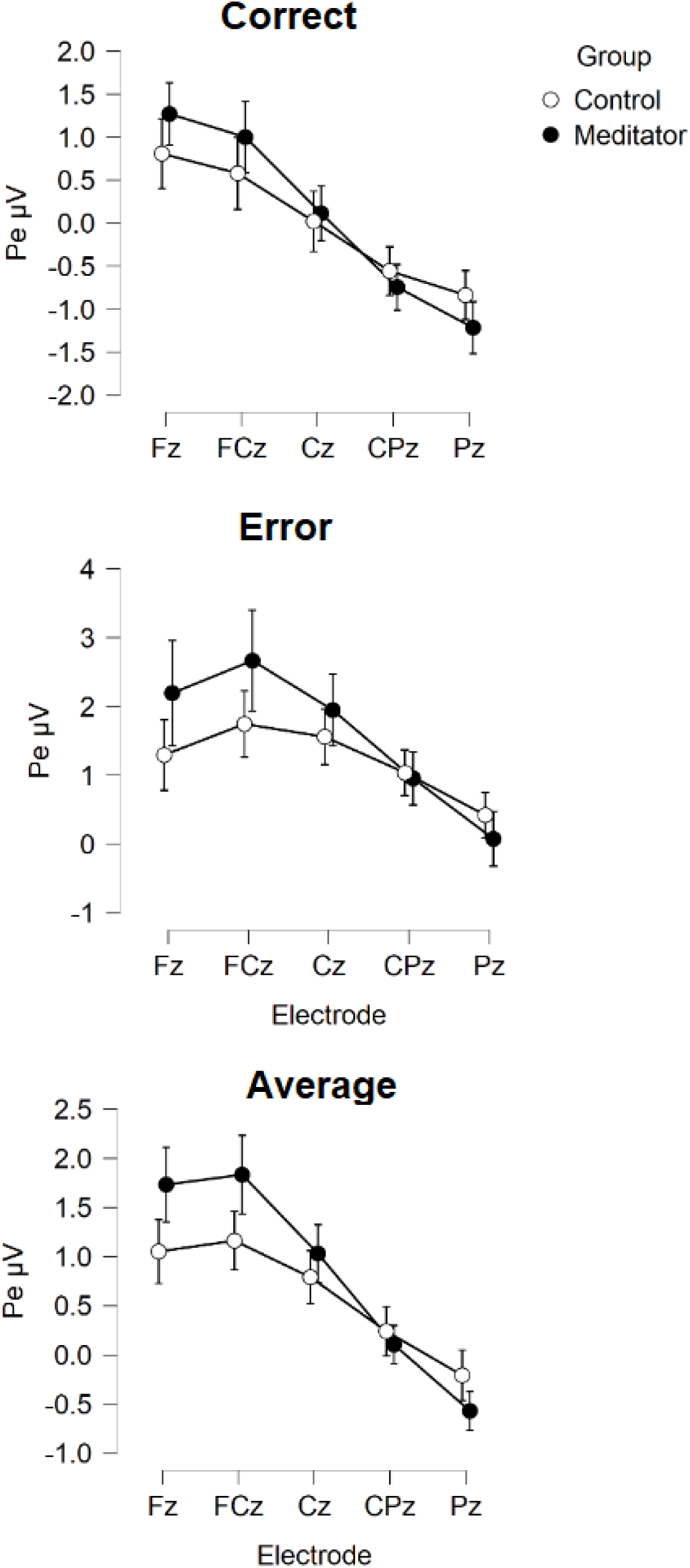
Activity during the Pe window for correct and error responses in meditators and controls from the novel dataset. Error bars reflect 95% confidence intervals.

When viewed in conjunction with the GFP results, this result suggests that the meditation group showed a pattern of stronger neural activation following both correct and error responses during the Pe window, while still showing the same distribution of neural activation (the meditation group was not activating different brain regions to the control group, just activating the same brain regions more strongly, which, given the dipolar nature of brain activity, resulted in stronger differentiation in voltage between frontal and posterior voltages). No other main effect of group or interaction involving group was significant (all *p* > 0.33, all BFexcl > 2.2, see table 3).

## Discussion

Previous research has provided inconsistent evidence for differences in EEG markers of error processing in experienced mindfulness meditators. We hypothesised that the experienced meditation group would show larger ERN and Pe responses. Our current results provide weak Bayesian and positive frequentist evidence supportive of larger Pe neural response strength in meditators for both correct and error responses, but no differences in the ERN. However, we note that the differences in overall Pe amplitude did not pass the test-specific multiple comparison controls in our primary analysis of the novel data. If we had focused our analysis restricted to errors (which show a larger neural response than correct responses), the differences in the Pe would have passed multiple comparison controls. Additionally, our single electrode analysis of the Pe did pass multiple comparison controls. Bayesian analyses of the single electrode comparisons and error only comparisons showed strong to extreme evidence in support of differences in the Pe between the groups, and the results were consistent when data were combined across both our current and previous datasets (N = 96, reported in the supplementary materials).

### The difference in Pe is not specific to errors

Although previous research has commonly focused on the error Pe, there was no significant interaction between group and response type. This suggests the difference between groups in the Pe was present following both correct and error responses. As such, a potential difference between meditators and controls in the Pe may reflect generic performance monitoring and awareness rather than error processing specifically. Previous research has suggested that the ERN to errors and correct responses might be a combination of both an error sensitive process, and another process related to response monitoring that is independent of whether a response is correct or not (Endrass et al., 2012). Research has also demonstrated that the Pe following correct trials is modulated by the speed of responses when participants are instructed to respond quickly, and that correct Pe amplitudes are related to confidence in the accuracy of a response, suggesting correct Pe amplitudes are still related to response monitoring (Boldt & Yeung, 2015, Valt & Stürmer). As such, it may be that our tentative difference in Pe amplitudes across both correct and error responses in the mindfulness group indicates that the outcome independent component of the Pe is modulated by mindfulness, or that both the outcome independent component and the error sensitive process are modulated (but that the effect on the error sensitive process is not large enough for us to detect a significant interaction). This perhaps makes more sense than an effect of meditation that is specific to having made an error of commission, as there is no suggestion from mindfulness practice or the effects of mindfulness on attention, self-awareness, and executive function that the effects of mindfulness on performance monitoring would be specific to having made an error of commission, rather than performance monitoring in general. Indeed, within the predictive coding theory, the sensory input processed following a correct trial is still “expected prediction error” (Alexander & Brown, 2019).

### What does the difference in Pe amplitude mean?

Any interpretation of the potential difference in the Pe in meditators must be predicated on a functional interpretation of the Pe. It is likely that the processes underlying potential differences in the Pe are common across other neural activities, and as such, explanations for the current results should consider mechanisms that are in common across other neural changes from meditation. In this context, we note that perhaps the most powerful explanatory model of neural functions available is the predictive coding theory (Huang & Rao, 2011). As described in the introduction, this model views the brain as a Bayesian prediction generator, which processes information by updating its prior model of the sensorium based on new sensory evidence. While research has not yet applied the predictive coding theory to interpret the Pe, the Pe has been suggested to indicate a neural marker for the accumulated evidence participants have access to in deciding whether they have committed an error (Steinhauser & Yeung, 2010). In that context, a Pe of larger amplitude is compatible with a predictive coding explanation of the effect of meditation that suggests meditators show increased synaptic gain for the processing of prediction errors, facilitated by increased expected precision (in the form of neural gain) to sensory evidence (Lutz et al., 2019, Lakonen & Slagter, 2021, Manjalay et al., 2020, Verdonk et al., 2021). This is also consistent with predictive coding models that suggest mindfulness increases processing related to the “experiencing self” (which may be reflected by increased gain on bottom-up prediction error processing), and reduces processing related to the “conceptual self”, found higher in the cortical hierarchy, within regions like the DLPFC (Laukonnen & Slagter, 2021). Note also that our results did not indicate the effect was specific to errors, suggesting that meditators may show constantly enhanced processing of prediction error. It is also worth noting that our results neither support, nor provide evidence against hypotheses that mindfulness meditation reduces the formation of or precision of priors, or hypotheses that suggest mindfulness reduces the amplitude of prediction error processing.

An enhanced Pe is also aligned with functional magnetic resonance imaging research into the effects of meditation, as the Pe is thought to be generated by the cingulate cortex and insula, both of which show increased activity as a result of meditation practice (Tomasino et al., 2013, Boccia et al., 2015, Fox et al., 2016). It is also worth noting that the difference in the Pe might overlap with our previous research using different tasks. This research demonstrated that meditators have enhanced frontal P3 amplitudes in a Go/Nogo task (BLINDED FOR REVIEW), and enhanced frontal positive activity during the same time window following probe stimuli in the Sternberg working memory task (BLINDED FOR REVIEW) and following correctly encoded stimuli in the Nback task (BLINDED FOR REVIEW). The Pe and P3 have been suggested to reflect similar underlying processes (a neural reaction to a stimulus in the case of the Go/Nogo and working memory tasks, and a neural reaction to participant response in the case of the Pe, regardless of whether it is correct or incorrect) (Ridderinkhof et al., 2009). These neural activity differences might indicate a common difference in neural responses to the environment, which may be characterised by stronger frontal positive voltages and more negative posterior voltages from approximately 280 to 380ms (similar to a P3a, or frontally distributed P3 activation), and may all reflect updating of prior beliefs by posterior evidence via the processing of prediction errors.

### Comparisons with previous research – the Pe

The aforementioned interpretations of the functional significance of a difference in the Pe in experienced meditators hinge on the current result being robustly replicated. The larger Pe amplitude in the meditation group is an effect in the opposite direction to Andreu et al., (2017), and conflicts with the null results of Teper and Inzlicht (2012) (studies which examined meditators with years of experience). It is also worth noting that our previous error processing study provided Bayesian statistical evidence for the null hypothesis of no difference between the groups. However, when the two datasets were combined, the analysis still provided strong evidence for the interaction between group and electrode (BFincl = 11.115). This suggests to us that the conflict between the results of our current study (showing a larger Pe amplitude) and previous studies (showing a smaller Pe amplitude or a null result for the Pe) might be of methodological origin.

One potential methodological explanation for the difference between the current study and our previous study may be the task used to measure error processing. The current study restricted analyses to EEG activity during the Go/Nogo task only (which has been shown to produce the most dependable error processing effects; Clayson et al., 2020), whereas the previous study included errors from the Go/Nogo task, the colour Stroop and the emotional Stroop task. However, we think this explanation is unlikely, as the Stroop task has been shown to still produce reasonably dependable error processing measures (Clayson et al., 2020). Another possibility is the data processing methods used. Our previous study used data pre-processing methods that we have since demonstrated to be less effective at cleaning artifacts compared to the method used in the current study (BLINDED FOR REVIEW). However, different EEG data cleaning approaches have been shown to produce only minor differences in study outcomes (Robbins et al., 2020), and these differences are still aligned in direction (Robbins et al., 2020, Bailey et al., 2022a, 2022b). As such, it seems unlikely that these explanations would produce such strong evidence for the null hypothesis if the results from our current study reflect the true result (we would expect more inconclusive results).

Following research from Alday (2021), we also think it is likely that the traditional subtraction baseline correction methods used in our previous study have a negative effect on error processing studies compared to a regression baseline correction. Indeed, when data from our previous study were cleaned in the same way as the current study, the previous study’s pattern was in the same direction as the current study (although the pattern was still not significant).

It should also be noted that the difference in Pe between groups was limited to the overall neural response strength, and we did not find differences in the distribution of neural activity. While our results on the surface may seem to align with previous research suggesting a larger Pe amplitude can be found after short mindfulness interventions (Lin et al., 2019, Smart and Segalowitz 2017), it is worth noting that post-hoc exploration of the difference in the electrodes of interest analysis indicated that the groups only differed at Fz and FCz, but the larger effect size was from a different pattern across the electrodes between the groups (with meditators showing a larger voltage differentiation between frontal and posterior electrodes). This contrasts with previous research, which has typically shown a group main effect difference between meditators and controls in an analysis of a single electrode or activity averaged across a small group of electrodes (Andreu et al., 2017, Lin et al., 2019, Smart & Segalowitz 2017). If we had restricted our analysis to more posterior electrodes (as some studies have), we would have concluded there was no difference in error processing in meditators. This is an advantage of the GFP and TANOVA analysis methods applied in the current study - they were able to reveal that the potential difference in the Pe is due to stronger activation of typically activated brain regions, rather than a different pattern of brain regions being activated in meditators (Koenig et al., 2011, Habermann et al., 2018).

In addition to these potential methodological explanations, we suspect there may be considerable variability in the effect of meditation on the Pe. Large samples of very experienced meditators may be the most likely to detect reliable effects. If future research is interested in resolving the conflict, we provide suggestions for a study designed for a robust resolution of the issue in the supplementary materials. Considering that experienced meditators are difficult to recruit in large numbers, the probable small effects of mindfulness on the Pe even after long-term practice, the lack of ability to draw conclusions around causation from cross-sectional research, and the low potential for clinical applicability from EEG research, our view is that determining whether there are differences between meditators and non-meditators in EEG measures of error processing should not be a high priority for future research. Instead, we think research using an experimental approach to determine whether more meditation leads to larger effect sizes for improved mental health would be more beneficial.

### Comparisons with previous research – the ERN

In contrast to the positive results for the Pe our results suggested there was no difference between meditators and controls in the ERN. When viewed in combination moderate Bayesian evidence against a difference between groups in our previous study (Bailey et al., 2019), the results indicate that long-term mindfulness meditation experience does not alter the ERN.

Given this seems to be the case, it is confusing to us that several studies have detected changes in the ERN after mindfulness practice using smaller sample sizes and less experienced meditators (Pozuelos et al., 2019, Fissler et al., 2017, Smart & Segolowitz 2017, Andreu et al., 2017, Teper & Inzlicht, 2012), sometimes after only a single session of meditation (Saunders et al., 2016). We would expect that changes resulting from mindfulness are likely to be detected following more extensive practice rather than less (Falcone & Jerram, 2018, Tomasino et al., 2013). It may be that brief mindfulness practice does affect ERN amplitudes, but that this change reverts to baseline after long periods of practice.

However, this explanation requires an additional assumption to explain the pattern of results across the study, and as such lacks parsimony and should be viewed with scepticism. A third potential explanation is that the wide range of potential analysis parameters available in EEG research could have provided analysis parameter selection biases towards positive results in previous research. These include the cleaning of muscle and blink artifacts from the raw EEG data, the choice of reference montage (Klawohn et al., 2020), the choice of baseline correction periods, the number of epochs for inclusion in the analysis, and the choice of electrodes and windows for analysis can all be varied by the experimenter and influence results. In particular, the choice of windows for analysis can be selected after inspection of group means, which has been demonstrated by simulations to inflate false positive rates (Kilner 2013), and a similar issue applies for the selection of electrodes for analysis. While we proposed that random variation in effect size across studies is a possible explanation for the inconsistency in results relating to the Pe, we think it is unlikely that this could explain two null results in independent datasets for the ERN. As such, given null results across two separate studies (both in highly experienced meditators and with reasonable sample sizes), we suggest that meditation is unlikely to affect the ERN.

Lastly, while we think it is likely that meditation does not affect the ERN, and think further research is needed before we can be confident in differences in the Pe, we note that null results do not mean that meditation does not affect neural activity. There is now robust evidence from meta-analyses that meditation does affect neural activity, particularly neural activity related to attention, self-regulation and interoception (Tomasino et al., 2013, Boccia et al., 2015, Fox et al., 2016). As such, the current findings provide subtlety to these findings, suggesting that while previous research has indicated meditation is likely to affect neural activity related to specific processes, it is unlikely that the processes underlying the ERN are altered by meditation.

### Limitations and Future Directions

While our study reflects the largest sample of experienced meditators collected to date, our primary comparisons were unfortunately underpowered to provide a conclusive answer regarding the Pe, so our exploratory analyses were the only analyses able to provide support for a larger Pe in meditators. Additionally, the meditation group in our study was compared against healthy control non-meditators (in contrast to a group who had also undergone a practice of some kind with equivalent intensity and duration to meditation training, for example athletes in the case of Andreu et al., 2017). The inclusion of an active control group would have been beneficial for controlling for non-specific factors that might have influenced our findings related to the Pe. However, the lack of control for these factors does not affect our conclusion of a null result with regards to the ERN (our results indicate that neither the meditation practice nor the other uncontrolled factors affected the ERN). Similarly, the cross-sectional design is a limiting factor when considering causation. However, it is worth recognizing how difficult it would be to provide good evidence with a longitudinal study – considering that despite the amount of meditation practice undertaken by our participants, our study did not even provide conclusive evidence for a cross-sectional difference in the Pe, and a longitudinal study with equivalent meditation experience and sample size would be almost impossible to practically implement.

Another potential limitation is that our decision to obtain a larger dataset by combining our previous data with the current data was not planned – and was implemented in order to explore why our results conflicted. However, to prevent any potential experimenter bias in this process from implementing our results, our inclusion of participants was based only on selection of individuals who provided enough error response epochs for analysis from the Go/Nogo task in our previous dataset, and all participants providing enough epochs from the earlier study were included. Additionally, because our previous study showed no differences between meditators and controls, an a priori assumption would be that combining the two datasets would have biased our results towards the null result rather than to strengthen the difference we found in the Pe in our novel dataset. As such, we believe the combined dataset is valuable in providing evidence that the Pe difference might reflect a real result.

Lastly, it is also worth noting that the study aim was to resolve the conflict between the null result from our previous study (BLINDED FOR REVIEW) and the research from a number of other groups. As such, we performed several exploratory analyses to ensure the lead author’s expectation for a null result did not bias our conclusions. This means that it is possible that the difference in the Pe is simply a false positive produced by repeated statistical tests. To address this, we did implement experiment-wise multiple comparison controls.

However, it is also possible that the number of multiple comparison controls implemented reduced our power to detect a significant effect in our primary analysis. We recommend that future research focus specifically on a single independent samples t-test comparing the GFP of the Pe between a group of long term meditators and a control group in order to maximise the chance of detecting a significant effect, while not inflating potential false positive results due to multiple comparisons.

