## Supplemental Materials for "Meditators probably show increased behaviour-monitoring related neural activity"

**Supplementary materials**

**Additional Background**

With regards to the previous literature examining error processing in mindfulness, three interventional studies have shown increased ERN amplitudes in the mindfulness group (Fissler et al. 2017, Pozuelos et al. 2019), or an increased difference between the ERN to correct versus error responses, following either single session mindfulness inductions, or a mindfulness intervention lasting many weeks (Saunders et al. 2016, although only for a “mindfulness of emotions” condition, not “mindfulness of thoughts”). However, one study examining participants with attention deficit hyperactivity disorder (ADHD) showed the opposite effect, i.e., reduced ERN amplitudes after mindfulness practice (Schoenberg et al. 2014); while four additional studies in healthy controls have shown no differences between mindfulness and control groups on ERN amplitude (Eichel et al. 2020, Larson et al. 2013, Lin et al. 2019, Rodeback et al. 2020). Results for the Pe are similarly inconsistent. Two studies reported an increase in Pe amplitude (Lin et al. 2019, Rodeback et al. 2020, although their “active” condition was a stress induction paradigm, and the mindfulness condition was viewed as the control), and one reported a trend towards an amplitude increase (Smart and Segalowitz, 2017). One study reported no difference (Bing-Canar et al. 2016), two studies reported an increase in Pe amplitude in the mindfulness group, but also an increase in the waitlist or relaxation control groups (Schoenberg et al. 2014, Eichel and Stahl, 2020), and one study reported a reduced Pe amplitude in the mindfulness group (Larson et al. 2013).

In addition to the four theories about how mindfulness might affect the parameters of the predictive coding theory provided in the introduction, we summarize a further four theories as follows: Mindfulness practice might 5) decrease the association between posteriors, prediction error, and active inference as a result of observing sensations (posteriors and prediction errors) without making active inferences to resolve prediction errors (sitting still and simply observing the sensation) (Deane et al. 2020); 6) increase the endogenous control of goal-related priors, which is proposed to reduce the unpleasant experience of prediction errors in favour of autonomic control over the predictive coding system (Deane et al. 2020), 7) increase precision weighting on hierarchically lower perceptual processes (for example the breath), which reduces the formation or elaboration of more abstract priors that are associated with more distant timepoints from the current moment, which has the effect of reducing conceptual processing (Laukkonen and Slagter, 2021); and lastly, 8) make priors more epistemically accurate and concurrently less precise, by adding the concept of “temporary” to the priors, through maintenance of the prior that all sensations (posterior evidence) are temporary (Laukkonen and Slagter, 2021).

**EEG processing methods that can result in false positives**

Importantly, we noted that previous research analysed the ERN and Pe using values obtained from specific electrodes and time windows of interest. ERPs are typically dipolar (including the ERN and Pe), such that negative voltages occur at some electrodes concurrently with positive voltages at electrodes on the other side of the dipole (for example, the Pe typically shows positive voltages at fronto-central electrodes and negative voltages at posterior electrodes). As such, the choice of electrodes for analysis has the potential reverse a detected effect within the same dataset. For example, if a mindfulness group generated a larger amplitude Pe, and one study measured the Pe at FCz (where positive voltages would be detected) while another measured it at Pz (where negative voltages would be detected), the two studies might show the opposite effect. Additionally, applying an arbitrary selection of a time window for analyses can inflate the potential for false positive results (a positive finding in the absence of actual differences between groups) (Kilner, 2013). To reduce the potential that arbitrary electrode or time window selection for examination of the ERN and Pe affected the results, we used advanced EEG statistical methods that allow for analysis of all electrodes across the entire epoch time window (while using data driven controls for multiple comparisons). This reduced the potential for false positives from window selection biases, as well as reducing the potential that we might miss significant results that are outside selected windows of interest.

Lastly, in addition to the issues related to the analysis of ERPs, the methods used to apply baseline correction to EEG data have recently been shown to pose potentially serious issues for the conclusions we draw from ERP research. Baseline correction is commonly applied in ERP research to reduce the impact of slow drift in the voltages of EEG electrodes on ERPs (caused by non-neural artifacts such as changing skin impedance or poor electrode connection to the scalp). This slow drift can mean that at the start of an EEG window of interest, voltages are >20 μV away from the 0 point. The neural response to the stimuli (the ERP) is then overlaid onto this drifted data (and ERPs are typically 2-10 μV in size). Without baseline correction, the signal to noise ratio of an ERP can be severely reduced, increasing the probability of null results when the alternative hypothesis is true. This drift can also produce false positive results if one group shows more drift than another. However, while baseline correction is important, typical baseline corrections simply subtract the averaged pre-stimulus or pre-response voltage from the active period (for example, in our previous research we subtracted the averaged EEG activity from -400 to -100ms before error responses from the EEG activity following the error). Recently, Alday (2019) demonstrated that this subtraction baseline correction method transposes a mirror image of the distribution of activity during the baseline period onto the active period, while ironically also decreasing the signal to noise ratio. Essentially, if EEG activity in the baseline period differs between groups, this difference baseline subtraction transposes the difference to the active period, and thus researchers might conclude that the active period differs even when it does not. This is a risk generally for ERP research, but particularly so for EEG measures of error processing, where the baseline period usually overlaps with ERPs timelocked to the stimuli. For example, in our Go/Nogo task, the EEG data within the baseline period of error response locked epochs displayed a topography typical of the N2 ERP in response to a Nogo stimuli (see the supplementary materials for a visualisation). Our previous research has shown that meditators differ from controls in stimulus locked ERPs (BLINDED FOR REVIEW), including in both the Go/Nogo task (BLINDED FOR REVIEW) and Stroop task (BLINDED FOR REVIEW), which are commonly used to examine neural activity related to error processing. As such, we implemented baseline correction by applying a regression and rejecting the variance contributed by the baseline period (which is the method recommended by Alday, 2019). This ensured our conclusions were not related to potential differences in the baseline period.

**Methods**

Note that the definition of mindfulness for the current study could include participants who practice both focused attention (FA) meditation and open monitoring (OM) meditation.

In the Go/Nogo task, for Go trials, participants were instructed to respond to stimuli using both index fingers to press separate buttons simultaneously. For Nogo trials, participants were instructed to withhold making a response. Participants completed a short practice round before the first block, and each block where the stimulus-response pairing changed. The second practice was included to prevent extra errors and reduce impact of task switching on ERN/Pe amplitudes.

To assess whether the number of accepted epochs from each response type for meditators and controls differed, a repeated measure ANOVA was conducted for the included number of epochs. Independent samples t-tests were used to compare the demographic and self-report variables between groups. Gender was compared using a Chi-squared test.

Because this study reports data only from a subset of participants who completed the task, behavioural data is not reflective of comparisons between groups, so is only reported as means rather than with statistical comparisons (which were not of interest in the current study). Full behavioural performance analyses will be reported in Bailey et al. (in preparation), which will examine neural activity related to response inhibition following stimulus locked neural activity in the Go/Nogo task.

For the number of epochs included from each participant, no main effect of group or interaction between group (meditators/controls) and task response (correct/error) was found (*p* > .2 and BF01 = 3.41 for the model including group and response * group, see Table 2).

**EEG data cleaning**

Continuous data were filtered with a fourth order Butterworth filter from 0.1 to 80Hz with a second order Butterworth band stop filter between 47 and 53Hz (to address noise from the electrical supply). Bad channels were detected and rejected using PREP (Bigdeley-Shamlo et al. 2015), then remaining channels were rejected if they showed extreme outlying data for more than 5% of the total recording (extreme data was defined as voltage shifts over a 1 second period >20 median absolute deviations [MAD] [with blink periods automatically detected and a threshold of >8MAD applied separately to 1 second periods containing blinks and to periods not containing blinks], a distribution of voltages within a 1s period that was more than 8SD from the mean or showed more than 8SD from the kurtosis mean, showed an absolute voltage of more than 500μV in a 1s period, or a frequency/power slope < -4 (suggesting drift without neural activity). Channels containing more than 5% of the data affected by extreme artifacts were rejected. If more than 20% of the channels were marked for rejection, the channels were ranked as to the amount of data affected by extreme artifacts, and only the top 20% were rejected. Next, channels with frequency/power slopes > -0.59 for more than 5% of the 1 second periods were rejected (suggesting muscle activity), again with channels ranked as to the extent of muscle affected periods, and only the most extreme 20% rejected if more than 20% were allocated for rejection. Following this, periods that still showed extreme data (as per the previously described definitions) were rejected.

The next step created masks of the remaining artifacts and submitted these masks to a sequential Multiple Wiener Filter (MWF) cleaning process (described by Somer et al. 2018). The first mask detected muscle activity, with frequency/power slopes > -0.59, and all 1 second periods (with 500ms overlaps) assessed for these slopes and marked as an artifact for up to 50% of the data (if more than 50% of the data exceeded the threshold, the cumulative amount that the slope exceeded the threshold across all electrodes was ranked across all epochs, and the most severe 50% only were marked). Marked artifact periods shorter than 1200ms had buffer time added to pad the period to 1200ms, in order to ensure sub-threshold artifacts just outside the 1s periods were captured in the MWF cleaning process. The second mask detected blinks, first filtering data with a 4^th^ order butterworth filter from 1 to 25Hz, then averaging data across blink affected electrodes (F3, F1, Fz, F2, F3, AF3, AF4, FP1, FPz, FP2), determining the 75^th^ percentile of voltage amplitude across all datapoints, adding 3 x the interquartile range to this value, and marking blinks at the voltage peak within the period that exceeds this threshold. 800ms periods surrounding each marked blink were marked in the artifact mask for the MWF cleaning. Third, periods showing drift (frequency/power slopes < -4) and horizontal eye movements (defined as concurrent shifts in voltage of >2MAD from the median in the opposite direction in the lateral electrodes on each side of the head for >25ms in total out of any 50ms period) were marked as artifacts for the MWF cleaning. 200ms of buffer was added to each horizontal eye movement artefact to ensure sub-threshold artifacts just outside marked periods were captured in the MWF cleaning process, and all artifact marked periods were extended to 1200ms if they were shorter than this duration. Prior to the MWF cleaning, artefact or clean data periods in the mask were marked as “exclude from both the artifact and clean masks” if shorter than 1200ms, and the first and last 5 seconds of the data file was marked similarly. A delay period of 8ms was implemented in the MWF cleaning, so that the artifact and clean data templates accounted for both spatial and temporal information (shown to enhance MWF cleaning performance).

After the three MWF cleanings were applied, infomax independent component analysis was performed using cudaICA, and ICLabel was used to identify artifactual components. These components were reduced using wavelet-enhanced ICA. Performing the MWF cleaning prior to the ICA provides the advantage of good ICA performance without having to high pass filter at 1Hz (which has been shown to reduce the sensitivity of ERP paradigms).

**ERA Toolbox Dependability Analysis**

The ERA toolbox provides a measure of dependability, which is a generalisation statistics metric that is analogous to measures of internal consistency, with the additional benefit of being able to examine the reliability of multiple data aspects within the same analysis. Dependability values range from 0 to 1, with 1 being maximum dependability. The ERA Toolbox was not designed for use on the topography of ERPs (as per the TANOVA in our study), so we extracted the GFP of the ERN averaged across the 50 to 150ms time window and Pe from 200 to 400ms post response, as well as the mean amplitude from electrodes where the ERN and Pe were maximum (Fz, FCz, FC1, FC2, Cz) and submitted these values from each trial from each participant to the dependability analysis. The overall dependability measure obtained for the 54 participants included in our combined dataset was 0.90 [0.85 to 0.93] for the correct trials Pe and 0.94 [0.92 0.96] for the error trial Pe GFP, suggesting our analysis produced dependable results. For the dependability analysis averaged across the electrodes showing the Pe maximum (Fz, FCz, FC1, FC2, Cz), we obtained dependability values of 0.90 [0.85 to 0.93] for the correct trials Pe and 0.94 [0.92 0.96] for the error trial Pe. The GFP dependability metrics for the ERN were 0.92 [0.89 0.95] for both the error and correct trials. The electrode of interest dependability for the ERN was 0.92 [0.89 0.95] for correct trials and 0.92 [0.89 0.95] for error trials.

**Electrodes of Interest Analysis**

In addition to the whole scalp Randomisation Graphical User Interface (RAGU) analysis conducted to test our primary hypotheses, a traditional electrodes of interest analysis was conducted. Activity occurring during the ERN window (defined as activity from 50 to 150ms following the response) and Pe window (defined as activity 200 to 400ms following the response) was averaged at midline electrodes Fz, FCz, Cz, CPz and Pz. Averaged voltage within the ERN and Pe windows were calculated for each participant for both error and correct responses. Frequentists analyses of single electrode data and Bayesian analyses were performed using JASP 0.13 (Love et al. 2019). Repeated measures ANOVAs were used to conduct a 2 group (controls vs meditators) x 2 trial type (correct vs error) x 5 electrode (Fz, FCz, Cz, CPz, Pz) comparison for the ERN and Pe separately.

Greenhouse-Geisser corrections were applied to control for violations of sphericity where necessary. Where interactions were significant, post-hoc t-tests were conducted to explore differences between the specific group/condition effects driving the interaction, controlled for multiple comparisons within the omnibus statistical test within each specific ANOVA using the holm method across all t-tests for all group/condition effects involved in the significant interaction. Bayesian repeated measures ANOVA analyses were used to determine the likelihood of the null hypothesis in contrast to the alternative hypothesis. For these analyses, we performed the suggested comparison between models containing a hypothesised effect to equivalent models stripped of the effect, to determine the likelihood of the alternative or null hypothesis for the specific main effect or interaction effect of interest.

For the electrodes of interest analyses, a total of 28 values were winsorised to the 75/25% +/- 1.5*IQR boundary, with a maximum of five values winsorised from any particular condition. Winsorised data for the novel dataset electrodes of interest analyses: 2 from correctFzERN, 2 from correctFCzERN, 1 from CorrectPzERN, 5 from the ErrorFzERN, 2 from the ErrorFCzERN, 2 from the ErrorCzERN, 3 from the ErrorCPzERN, 2 from the ErrorPzERN, 2 from correctCzPe, 1 from CorrectPzPe, 2 from ErrorFzPe, 2 from errorFCzERN, 1 from errorCzPe, and 1 from errorPzPe.

Levene’s test indicated no violations were present in the assumption of equality of variances, and Shapiro-Wilk tests for each variable indicated normality (all *p* > 0.10) with the exception of the correct response ERN at Cz for the meditator group and the correct response ERN at Pz for the control group (p = 0.025 and p = 0.009 respectively). However, parametric statistical tests showed no significant differences involving group for the ERN, and the RAGU analyses (which are robust against violations of the assumptions of parametric statistics) showed no differences, so the null result was assumed and no correction was applied.

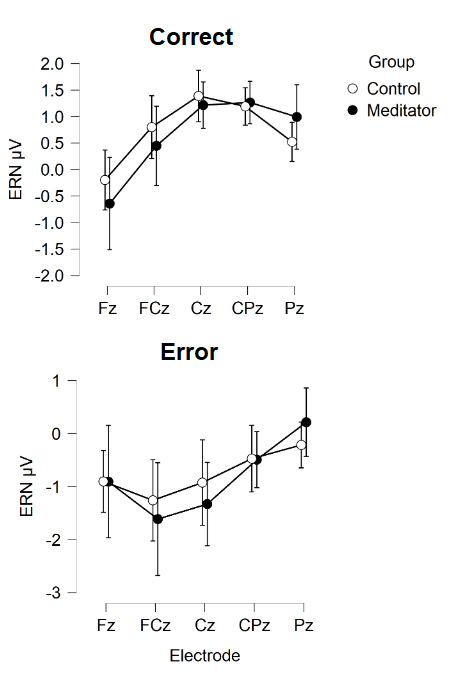

Figure. S1 Activity during the ERN window for correct and error responses in meditators and controls from the current study’s dataset. Error bars reflect 95% confidence intervals.

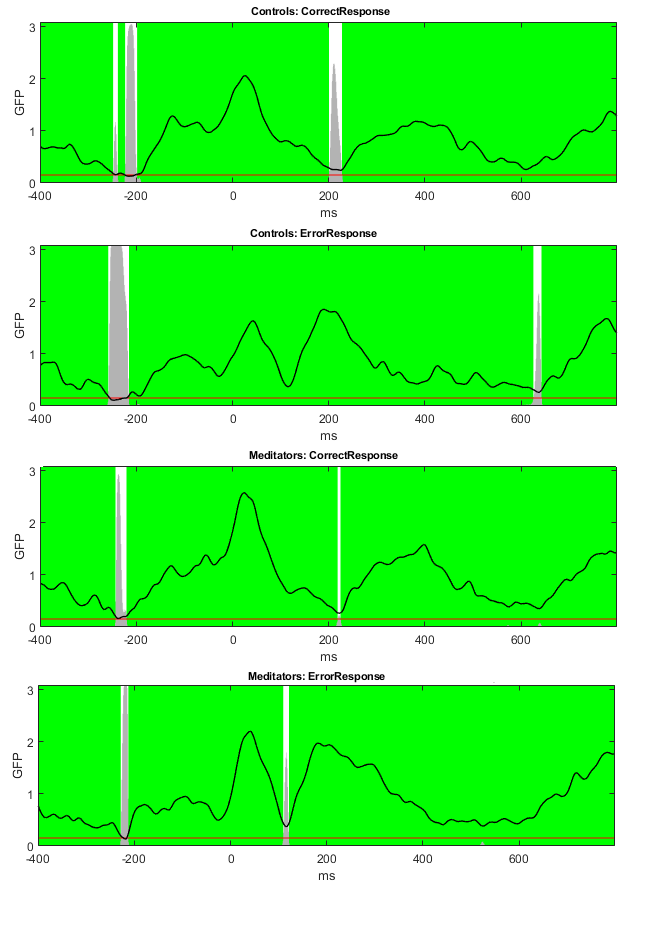

Figure S2. TCT results.

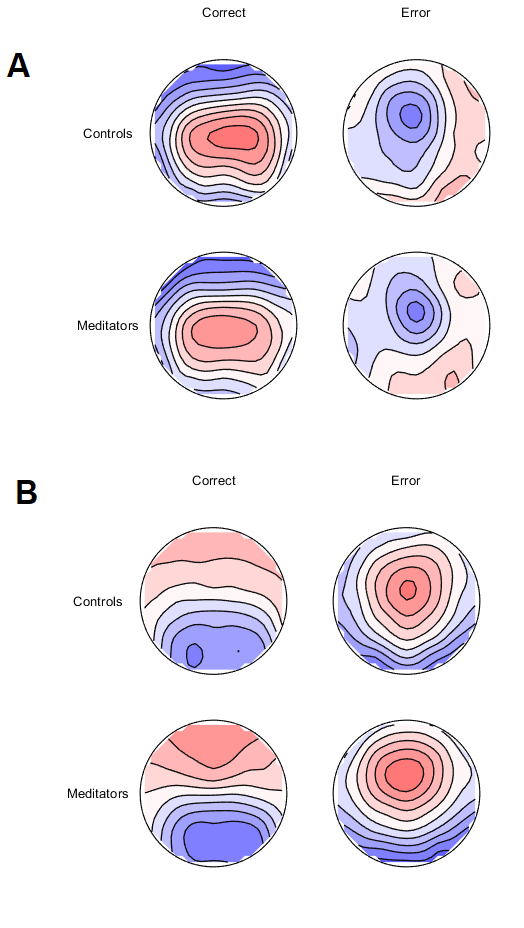

Figure S3. The distribution of activity from the current study’s dataset, averaged A: within the ERN window (50 to 150ms) and B: within the Pe time window (200 to 400ms) following the response.

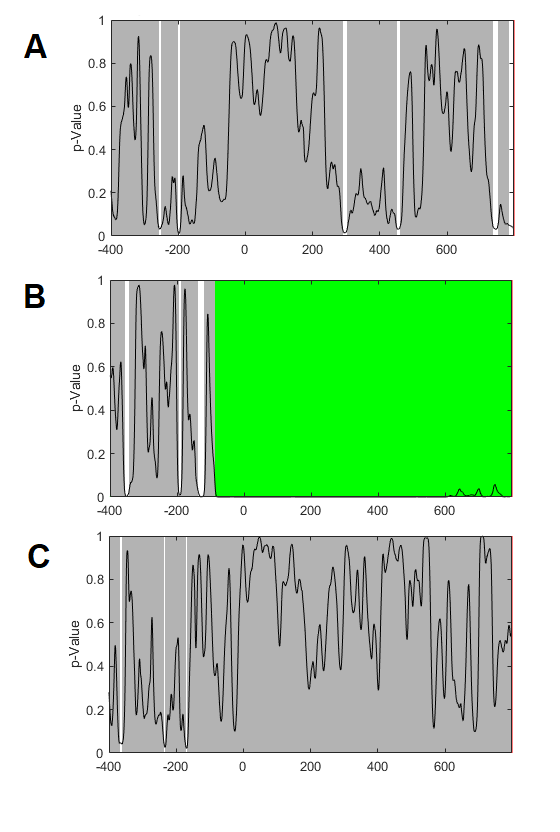

Figure S4. p-graphs for the TANOVA comparison in the current study’s dataset main effect of group (A), main effect of condition (B), and group by condition interaction (C).

**Re-analysis of our previous dataset**

In order to determine the potential cause of the conflict in results between our previous study (N = 42) and our current study, we re-processed the data from our previous study using our new data cleaning approach, and performed a regression baseline correction on each error related epoch (as per our current methods), in contrast to the traditional baseline subtraction method (which we conducted in our previous study). Averaged across the Pe window of interest (200 to 400ms), there was no significant difference between the groups (p = 0.176, np^2^ = 0.0576, BF10 = 0.658). However, in alignment with the results of our current study, the meditator group showed a non-significant but larger error related Pe GFP. Data can be visualised in Figure S8. No significant differences were present for the single electrode analysis either, with the group main effect showing no significant difference F(1,40) = 0.614, p = 0.438, np^2^ = 0.015 and the group by electrode interaction showing no significant effect F(1,40) = 0.951, p = 0.378, np^2^ = 0.023. However, the pattern was again similar to the pattern in our current study (Figure S9).

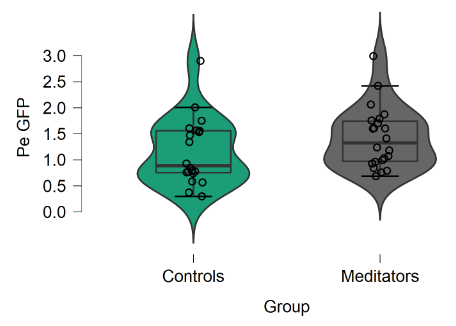

Figure S5. The Pe GFP from our old dataset.

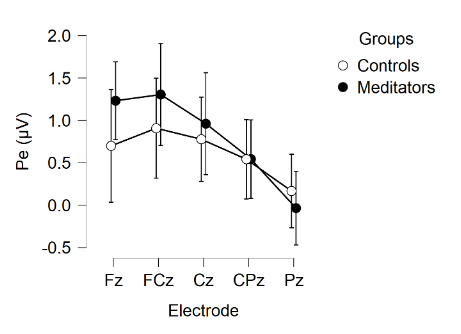

Figure S6. Pe amplitude from a single electrode analysis of our old dataset after using our new cleaning method on the data and using the baseline regression method.

**Combined dataset**

We combined our old and new dataset together (N = 96) to obtain a more complete understanding of any potential differences in experienced meditators in the Pe marker of error processing, provided by a larger and more generalisable sample. Similar to the results of our current study, averaged across the Pe window of interest (200 to 400ms), the meditator group showed significantly larger error related Pe GFP (p = 0.019, np^2^ = 0.0576, BF10 = 2.607, or BF10 = 5.147 for a one-sided test that meditators would show larger Pe GFP, given the sample in our current study suggested this would be the pattern). Data can be visualised in Figure S10. Also similar to the results of our current study, a significant interaction was present for the single electrode analysis, F(1,94) = 4.291, p = 0.025, np^2^ = 0.044, BFincl = 11.115. The pattern was again similar to the pattern in our current study (Figure S11). The group main effect showed no significant difference F(1,94) = 2.205, p = 0.141, np^2^ = 0.023, BFexcl = 2.031.

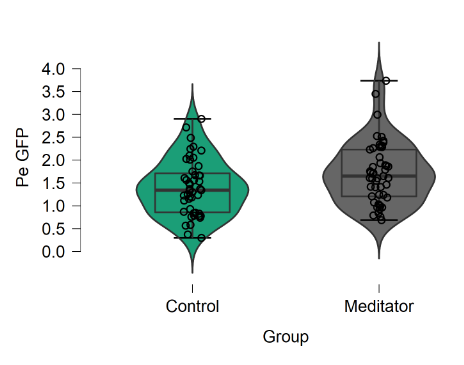

Figure S7. The Pe GFP from the combined dataset.

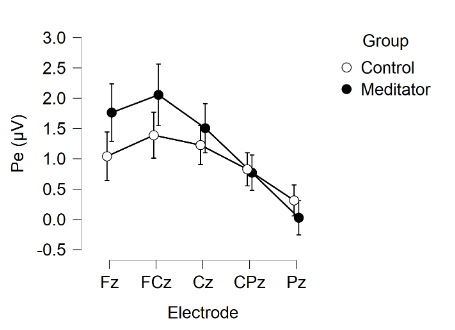

Figure S8. The single electrode analysis from the combined dataset.

| Whole combined dataset | Meditators,  *M*(*SD*) (*n* = 47) | Controls,  *M*(*SD*) (*n* = 49) | Statistics |
| --- | --- | --- | --- |
| Age | 32.19 (13.01) | 35.10 (12.09) | t(94) = -1.136, p = 0.259 |
| Gender (M/F) | 30/19 | 24/23 | Chi-square = 1.465, *p* = 0.226 |
| Years of education | 16.51 (2.97) | 16.42 (2.75) | t(94) = 0.151, p = 0.880 |
| Pe GFP | 1.717 (0.674) | 1.402 (0.610) | p = 0.019, np^2^ = 0.0576, BF10 = 2.607 |
| Fz | 1.762 (1.65) | 1.041 (1.37) | Group main effect: |
| FCz | 2.055 (1.77) | 1.389 (1.29) | p = 0.141, np^2^ = 0.023, BFexcl = 2.031 |
| Cz | 1.505 (1.41) | 1.226 (1.10) | Group x electrode |
| CPz | 0.770 (1.01) | 0.826 (0.93) | p = 0.025, np^2^ = 0.044, BFincl = 11.115 |
| Pz | 0.026 (0.98) | 0.313 (0.87) |  |

Table S1. Participant demographic, Pe GFP and single electrode data for the combined dataset.

Finally, because the two larger groups from the combined dataset were not directly matched in age (although they did not significantly differ in age), we conducted a final age matched analysis by randomly selection participants to exclude from each group so that every participant was directly age matched to within 3 years of one unique participant in the other group.

To do this, we programmed a MATLAB script to randomly shuffle the control participants, then select without replacement from the meditation group a list of unique participants who were within 3 years of age from each single participant in the control group (using the randsample function, selecting from all participants that were within 3 years of age of each single control participant, for each control participant). Once this group of meditators was obtained, we reversed the process, and randomly selected a control participant matched to within 3 years of age of each meditator participant (from the pool of control participants who were within 3 years of age for each meditator participant separately). We set the function to repeat until the number of unique participants in each group was equivalent to the number of participants in the overall combined group, or until the process had run through 10,000 iterations. After 10,000 iterations, we reduced the number of participants required in each group by 1, and ran the 10,000 iterations again. We repeated this process until the script met the conditions of having run through 10,000 iterations of attempts at randomly selection larger group sizes without obtaining those group sizes, and a maximum possible group size from 10,000 randomisations was selected.

The final age matched groups were 38 meditators and 38 controls long, and represented two directly age matched and randomly sampled groups of participants. As expected after the randomisation process, ages were well matched (meditator mean = 36.42, SD = 12.73, control mean = 34.74, SD = 13.24, p = 0.574, BF01 = 3.666)

When the error GFP was compared between these groups, results showed that meditators had a significantly larger GFP than controls from 249 to 318ms, which passed global duration controls (51ms), and meditators also showed significantly larger GFP than controls in another period immediately after this from 329 to 369ms (which did not pass global duration controls). Averaged across this period, the result was significant (p = 0.0114, partial eta squared = 0.0830, BF10 = 3.900). Averaged across the Pe window of interest (200 to 400ms), the result was also significant (p = 0.0330, partial eta squared = 0.0585, BF10 = 1.678). Including age as a covariate in the analysis also provided a significant result for both the specific window showing significance in the RAGU comparison, F(1,73) = 7.001, p = 0.01, partial eta squared = 0.088, BFincl = 4.078). Similarly, including age as a covariate in the analysis also provided a significant result for both the traditional Pe window (200 to 400ms), F(1,73) = 4.941, p = 0.029, partial eta squared = 0.063, BFincl = 1.751). These results demonstrate that the larger Pe GFP in meditators is robust against the potential confound of age.

**Study Design for Optimal Detection of a Difference in the Pe from Mindfulness Meditation**

Since our results suggest the error related Pe showed the strongest potential for detection of a difference, future research should perform a single comparison of the error related Pe. We suggest future research should extrapolate from the effect size for our significant Pe window from our combined dataset (Cohen’s d = 0.489). Given the inconsistency across the research, we suggest it would be most valuable to aim for 0.95 power. Using a one tailed approach requires a sample size of 92 per group to obtain 0.95 power with an alpha level of 0.05 (or 53 each group for 0.80 power). Note also that we recommend the use of dependability (Clayson et al. 2020) to determine how many epochs from each participant for inclusion, and as a result it is likely that even up to 20% of the data may have to be excluded. We also highly recommend using the regression approach to baseline correction rather than the subtraction approach. The Go/Nogo task seems best suited for producing dependable results and making this task difficult (with short time delays between trials and a high proportion of Go trials) and contain many trials (>300 trials) is the best way to obtain enough trials for each participant. It is also likely that there will be variability from study to study in the exact Pe time window after an error response that shows the effect, so approaches that provide the flexibility to detect a window of effect that is not specified a priori would be an advantage (while using data drive multiple comparison controls).

**The distribution of activity in our baseline correction period showed N2-like activity**

A common baseline period selected for baseline correction of error-related ERPs is the -400 to -100ms period before the error response. Figure S12 depicts the topographies averaged within this -400 to -100ms period (without any baseline correction applied to the data). These images show how the subtraction method would transpose a topographical distribution similar to the typical N2 onto the active period. The N2 topography is in fact almost the exact mirror of the Pe. As such, using the subtraction baseline correction method would almost certainly modify the Pe amplitude, and given that differences have been found in the N2 in meditators, might affect the results of the comparison between our meditator group and control group (note that the topographies did differ between conditions and groups in this period, although not at the level of significance, p = 0.426).

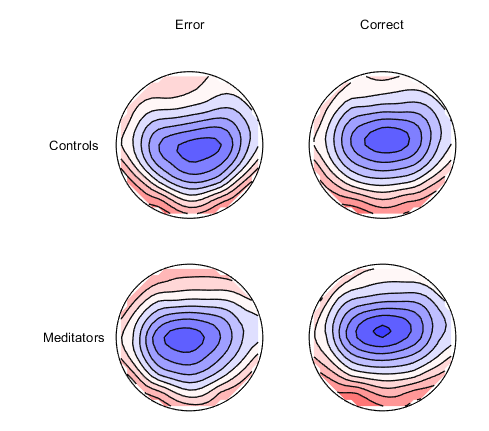

Figure S9. Topographical distribution of activity from each condition and group averaged within the -400 to -100ms baseline period (with no baseline correction applied to the data).
